# Inside Out: Functional Flexibility in Assigning the Body’s inside-outside

**DOI:** 10.1101/2025.09.10.675268

**Authors:** Jess Hartcher-O’Brien, Vincent Hayward, Malika Auvray

## Abstract

Tactile surface perception is often assumed to reflect a fixed boundary between body and world, with the skin providing a stable oriented surface that separates inside from outside. We show that this assumption fails in functionally specific ways. Using moving tactile stimuli on the hand, participants perceived stimulus direction on the fingertips as reversed when hand posture changed - reporting motion as if it originated from the opposite side of the skin, while responses on the palm were less consistent and less sensitive to posture. Congenitally blind participants exhibited stable, individualized surface assignments on the fingertips, demonstrating that vision is not required to construct oriented tactile surfaces. However, visual experience modulated posture effects: sighted individuals showed posture-dependent, skin-orientation assignments driven by motion direction, whereas congenitally blind individuals showed posture-invariant assignments determined by functional use. These findings indicate that tactile surface perception is not globally fixed but dynamically inferred according to functional demands; rather than enforcing a purely geometric body-world boundary, the brain flexibly assigns where the surface is and which side we occupy.

## Introduction

How does the brain distinguish the boundary between the body and the external world? A defining feature of physical objects, including our own body, is that they are volumetric solids with orientable surfaces that separate inside from outside (1). This boundary is not just a physical property but also a perceptual one: we interact with objects by perceiving their surfaces as having a consistent orientation, despite changes in our body’s surface orientation relative to objects (2).

In touch, this distinction implies that the skin should serve as a stable, oriented boundary, allowing the brain to assign sensations reliably to the “outside” surface of the body, even as our limbs change orientation during interaction and manipulation (e.g., (3–5)). We can thus think of the body as a unified volume, in which a consistent distinction between inner and outer surfaces is maintained throughout.

In geometric terms, a solid object can be described as a connected, continuous three-dimensional manifold embedded in 3D space (ℝ^3^). As such, being a solid ensures a stable inside-outside distinction, fundamental to how we experience objects as sided solids. Included in this category is our own body. In contrast, the brain could construct the skin as non-orientable surfaces such as the Moebius Strip, which lack this inside-outside distinction, having only one continuous side. Which geometric construct the brain enforces on the skin, the orientable-solid nature of 3D objects or the single-sided nature of a Moebius Strip, has not been thoroughly tested. However, tactile perception is known to respect surface orientation, as evidenced by studies of tactile reference frames. For example, a marker moving clockwise on the outer surface of an object, like the finger, would appear to move counterclockwise when viewed from the inside, demonstrating how perception is sensitive to surface orientation (Figure 1A). In tactile perception, analogous reversals would be expected if stimuli were applied to opposite sides of an oriented surface (6–9).

**Figure 1:**
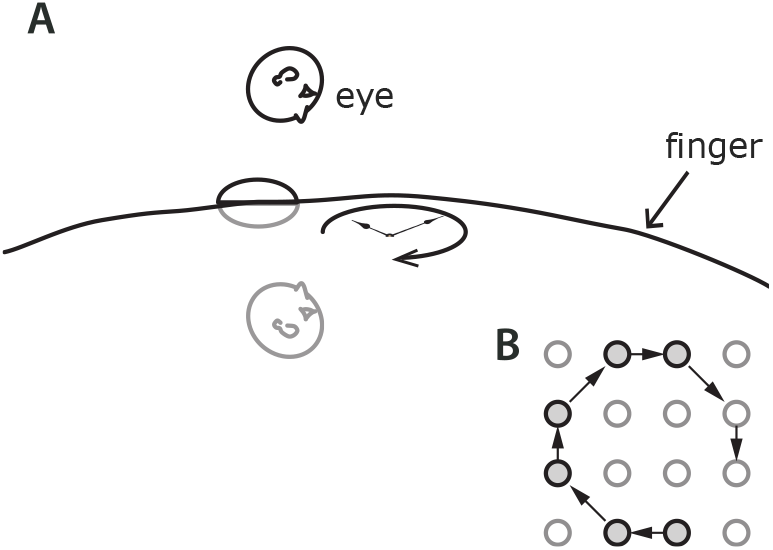
Surface orientation. (A) An observer sees, hears, or feels an object twirling in one direction on a surface, the finger. An observer “inside” the surface would see, hear, or feel the same object twirling in the opposite direction. (B) Clockwise pattern displayed by the tactile device.

Importantly, the skin of the hand is functionally heterogeneous: the palm provides a stabilizing contact surface, while the fingertips actively explore and manipulate objects from multiple orientations (10; 11). This functional distinction raises the possibility that surface orientation may be perceived differently across body parts. If so, it would suggest that tactile surface orientation is not fixed by geometry alone but is also shaped by the functional role of the body region.

We formalized this question through two competing hypotheses: First, the skin may be perceived as a surface with a globally fixed orientation - preserving the inside-outside distinction across postures and body parts. Alternatively, surface orientation might be assigned flexibly and locally, influenced by the function of each region of the skin.

To test these hypotheses, we applied tactile motion stimuli, that is stimuli rotating either clockwise or counter-clockwise, to the palm and fingertips across a range of hand postures (*Figure 1B*). If the skin is perceived as a stable, oriented surface, then the perceived direction of motion should remain consistent across postures. If perception is more flexible, we expect differences between the palm and fingertips: the palm, used for stabilization, may support stable but potentially weaker surface inferences, while the fingertip, with its exploratory role, may show flexible or even reversed surface assignment.

To determine whether visual experience contributes to the formation of these perceptual surface assignments, we also tested congenitally blind individuals. If visual input is necessary for building invariant surface representations, their performance should differ from sighted individuals. If not, this would suggest that functional roles and tactile experience alone are sufficient. It is often suggested that tactile spatial perception relies on calibration from vision early in life. As such the spatial assignment of tactile patterns would be expected to both differ from sighted to congenitally blind individuals and be more orientation-consistent for the visually impaired individuals.

Standard models of tactile spatial perception often rely on the notion of reference frames, in which sensations are localized by transforming their coordinates between distinct representational systems (e.g., body-centered, hand-centered or external frames)(8; 10). While widely applied across medical science and perceptual literature, this framework implicitly assumes that body parts function as modular scaffolds to which sensory signals must be remapped through coordinate transformations (12). Yet this approach may obscure more fundamental properties of body representation. In particular, it treats the body as a set of disconnected parts requiring spatial reconciliation, rather than as a unified perceptual volume.

Recent evidence from body-augmentation studies challenges the necessity of such referential constructs. For example, in a longitudinal training study, participants integrated a robotic sixth finger (D6) into the functional and neural architecture of the hand without any explicit spatial remapping (13). The D6 became co-represented with biological digits in the somatosensory cortex (S1), and was incorporated into coordinated motor behavior across the entire hand. Crucially, this integration occurred without competition for cortical territory or reference frame translation - suggesting that the sensorimotor system may operate within a unitary representational volume, capable of flexibly accommodating new effectors directly.

When applied to touch, this perspective suggests that the brain may not represent the body as separate parts with their own coordinate systems, but rather as one continuous surface within a single, unified space. From this viewpoint, understanding the direction of tactile inputs, like the way something moves across the skin, does not require translating between different spatial maps. Instead, it may rely on how each part of the skin is naturally shaped and used. This way of thinking better fits with how flexible the brain is in adapting to changes, and our results support this: the sense of direction on the skin is not determined by fixed geometry, but is built flexibly across the body.

Specifically, our results show that the brain does not impose a fixed inside-outside boundary on the skin’s surface. Instead, surface orientation is flexibly constructed - shaped by the functional role of each body part and influenced by sensory experience. This flexibility is especially pronounced at the fingertips, whose exploratory role permits a dynamic, context-sensitive tactile representation of the body. These findings challenge classical assumptions of the body as a fixed, volumetric solid in tactile perception.

## Results

To investigate how the brain assigns inside and outside to different regions of the body, we operationalised the notion of surface orientation on the volar surface of the hand, using a tactile device. The device displayed a randomised sequence of stimuli turning on the skin in clockwise and counterclockwise directions to the target location across the hand, see *Figure 1B*. The participants reported in which direction they felt the motion. Across three experiments, we tested whether the perceived direction of motion respects the geometric property of surface orientation and, critically, whether this assignment varies as a function of its location on the hand, which can be linked to differences in the hand’s functional role.

### Experiment 1: Surface assignment varies across the hand and reflects a functional role

Ten participants received stimuli at eight distinct locations across the volar side of the hand (*Figure 2*), ranging from the fingertips - primarily involved in active exploration - to more central palmar regions associated with support and stabilization. In this upright posture (hand pronated, fingers pointing upward), motion stimuli presented on the fingertip were reliably interpreted as rotating in a direction consistent with being felt from the “outer” surface of the hand. In contrast, motion applied to more central palmar locations was often reported in the opposite direction - suggesting a perceptual reversal, as if the stimuli were felt from the opposite (inner) side of the skin.

**Figure 2:**
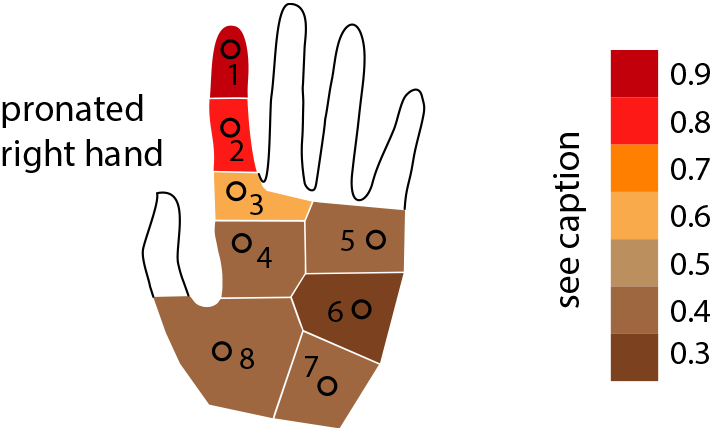
Experiment 1. Stimuli were applied to eight locations on the volar surface of the hand. Colour indicates the proportion of responses consistent with motion felt from inside the hand, reflecting perceptual reversal of surface orientation.

Critically, this transition from orientation-consistent to reversed percepts followed a gradient from fingertip to palm, aligning with the hand’s functional differentiation (11). If the brain treated the hand as a physical solid, responses should have preserved a consistent sense of surface orientation across locations. Instead, the pattern of perceptual reversals suggests that the representation of surface orientation is not fixed across the hand but flexibly assigned depending on function.

A mixed-effects logistic regression predicting the log-odds of reversed perception by location (with participant as a random effect) confirmed this behavioural pattern. The fingertip served as the reference and was consistently treated as part of an oriented solid. Adjacent location *02* did not differ significantly from the fingertip (*β* = 0.16, *p* = 0.63). However, responses from more central palm regions *03 to 08* were significantly more likely to indicate reversed surface assignment (all *ps <* 0.01), with the strongest effects at *06* and *07* (*β* = 2.45 and 2.16, respectively). These findings suggest that tactile perception of surface orientation is spatially heterogeneous, in a manner that reflect the functional role of each region rather than a fixed body-centered frame.

### Experiment 2: Perceived surface orientation depends on posture and function

A new cohort of participants (*N* = 15) received identical tactile motion stimuli to the fingertip or the palm across four hand postures: vertical /horizontal and pronated / supinated. If tactile perception consistently respects the geometric surface orientation of the skin, then the perceived direction of motion should remain constant. However, if perception reflects posture and function, then changes in hand orientation should systematically modulate the perceived direction. We collapsed responses across clockwise and counter-clockwise motion directions, as there was no significant difference in the proportion of direction-consistent responses across postures between the two directions for the fingertip (*F* (3, 84) = 0.082, *p* = 0.97) and the palm (*F* (3, 84) = 0.34, *p* = 0.80), respectively. The individual motion direction results can be seen in supplementary material *Figure S2*.

A two-way repeated measures ANOVA revealed a significant main effect of posture (*F* (4, 56) = 70, *p <* 0.001), but no main effect of region, indicating a similar overall pattern for both fingertip and palm. Critically, when the hand was flipped from a pronated to a supinated position, the perceived direction of motion on the fingertip reversed, indicating a loss of surface orientation consistency.

A significant interaction between posture and stimulated surface was also observed (*F* (4, 56) = 4.23, *p <* 0.05). Tukey’s post-hoc comparisons revealed robust differences in motion direction perception between fingertip and palm across all postures (*p <* 0.001). This supports the idea that the fingertip’s exploratory role involves a dynamic and posture-dependent assignment of surface orientation.

In contrast, responses on the palm showed weaker posture dependence, suggesting a less flexible or less functionally tuned surface assignment. These findings replicate and extend the functional dissociation observed in Experiment 1, demonstrating that the perceived orientation of tactile stimuli is shaped not only by body posture but also by the functional role of the stimulated region.

### Experiment 3: Surface assignment in blindness reveals the predominant role of function over vision

To identify the influence of long-term visual experience in addition to functional use, we tested eight congenitally blind individuals using the same design as Experiment 2. If surface orientation perception depends on visual calibration, blind participants should show degraded or absent orientation-invariant representations. Conversely, if functional engagement is sufficient to shape surface assignment, then consistent patterns should emerge.

On the fingertip, all blind participants exhibited posture-invariant responses - each showed a consistent perceptual interpretation across all orientations. However, the direction assigned, that is, what counted as clockwise, varied across individuals, indicating an idiosyncratic but stable sense of surface orientation. In contrast, responses on the palm were more consistent, and simultaneously less consistent across both postures and individuals, than sighted individuals.

**Figure 3:**
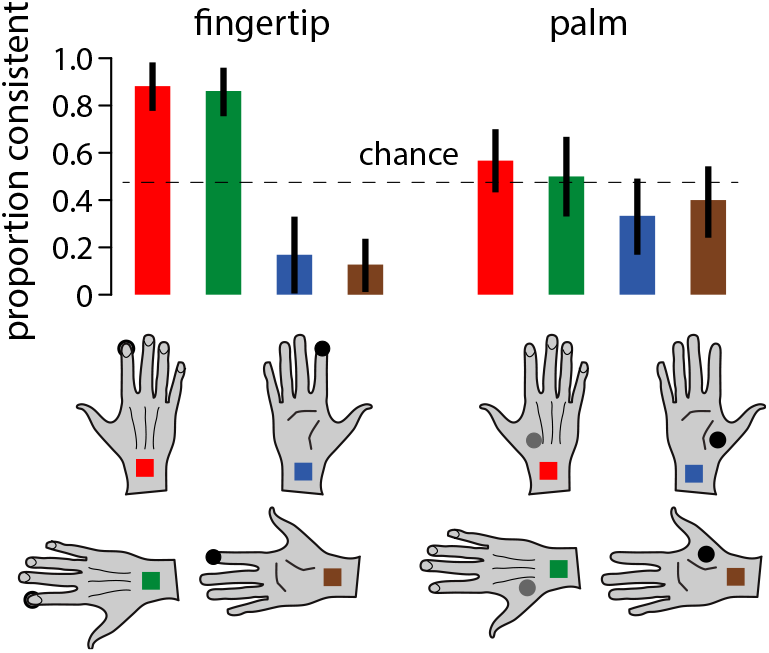
Experiment 2. Proportion of responses consistent with motion felt from the inner surface of the hand, by posture and location. Perceptual reversal was prominent on the fingertip and modulated by posture.

To quantify within-subject invariance, we conducted a nonparametric permutation test comparing the observed standard deviation across orientations to a null distribution generated by shuffling posture labels. The fingertip yielded a low mean standard deviation (*M* = 0.0640) and a permutation *p*-value of 1.000, indicating highly invariant responses across orientations. The palm produced higher variability (*M* = 0.0873), with a similar *p*-value of 1.000, indicating that any consistency could be explained by chance.

These results support the conclusion that visual experience matters in resolving inside-outside across posture. Given the consistency across both blind and sighted individuals in hand region-dependent behaviour, our findings further suggest that function is the primary driver of orientation-invariant surface perception. The fingertip’s consistent role in active exploration appears sufficient to develop a stable, individualized surface map - even in the absence of vision.

**Figure 4:**
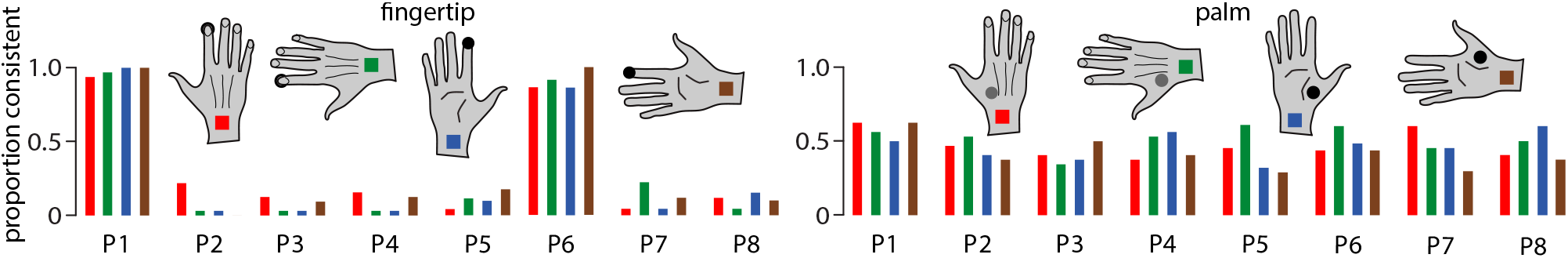
Experiment 3. Proportion of responses consistent with stimuli felt from the inner surface of the hand, by posture. Each line represents a blind participant. Stable, posture-invariant patterns were observed on the fingertip but not the palm.

## Discussion

Our findings challenge the long-standing assumption that the brain imposes a fixed, geometrically consistent, orientation on the skin’s surface (5; 13). Across three experiments, we show that motion direction, a marker of tactile surface orientation, is not veridically represented by the brain. Our results suggest that surface orientation is not globally preserved but flexibly constructed. In conjunction with visual experience, it is shaped by the posture and functional role of the stimulated region.

In Experiment 1, tactile motion on the fingertips which are specialized for active exploration, was perceived in a direction consistent with a stable, outward-facing surface. In contrast, motion on the palm, which is typically used for support, often yielded reversed interpretations, as if felt from the opposite side of the skin. This functional dissociation suggests that the brain does not apply a uniform inside-outside distinction across the body, but infers surface orientation based on behavioral relevance.

Experiment 2 revealed that posture further modulates surface assignment. When the hand was flipped, perceived motion direction on the fingertip reversed, demonstrating that tactile orientation is not treated as a fixed feature of the skin but is dynamically reassigned in context. Crucially, this remapping cannot be explained by differences in tactile acuity or standard coordinate transformations (9; 14), but reflects a functional strategy: perceptual stability is granted where consistent surface orientation supports interaction.

This principle echoes findings from haptic object recognition behaviour. For instance, Newell and colleagues showed that viewpoint-tolerance in touch is shaped by typical exploration paths. Our results suggest a similar mechanism within the body: tactile perception emphasizes exploratory configurations, constructing stable surface assignments where they are most behaviorally useful.

Experiment 3 provided further evidence for a functional origin of surface orientation. Congenitally blind participants showed consistent, posture-invariant responses on the fingertip with individualized mappings across participants. This suggests that visual input is not required for generating stable surface representations. Instead, tactile experience and functional use are sufficient to establish orientation, albeit in a personalized manner. These findings extend prior work on spatial reference frames in blindness (15), highlighting function - not modality - as the key organizing principle.

Real-world expertise supports this view. Braille readers, for example, show fluent motion interpretation across postures, reflecting a robust, posture-invariant surface assignment grounded in task demands. While cross-modal input may help standardize mappings across individuals (16), our results show it is not necessary for constructing stable, functionally tuned tactile orientation.

Together, these findings suggest that tactile surface orientation is not an inherent property of the skin but a dynamic perceptual construct, shaped by posture, sensory experience, and functional relevance. The fingertip consistently supports stable, context-sensitive assignments; the palm does not. This reframes the skin as a behavioral interface, not a passive sensory boundary.

Inherent in the functional perspective is a new lens from which to view classic graphesthesia phenomena (17–23; 7). In graphesthesia identical tactile letter / number traces are perceived differently depending on body location. Rather than invoking visual imagery or egocentric transformations alone, such effects may reflect region-specific surface assignments shaped by functional engagement and multisensory learning.

More broadly, our results contribute to ongoing debates about tactile reference frames (14; 24), showing that the brain prioritizes local, functionally relevant representations over global anatomical or external coordinates. This mirrors the flexibility observed in object-centered haptics and supports efficient multisensory integration for goal-directed action.

## Methods

### Participants

Five male and five female observers participated in Experiment 1 (age range: 19-35 years). Eight male and seven female participants took part in Experiment 2 (age range: 22-35 years). Experiment 3 included eight congenitally blind participants (four female, four male; age range: 26-35 years). All participants gave written, informed consent prior to the study, which was conducted in accordance with institutional ethics guidelines.

### Apparatus

The tactile stimuli were generated using a custom-built device consisting of two adjacent refreshable Braille cells forming a 4 × 4 pin array (25). Pins were activated sequentially to produce an apparent motion stimulus rotating either clockwise or counter-clockwise at a speed of 20 *mm/s*. Each stimulus lasted approximately one second. The apparatus was mounted on a stand, and participants rested their elbow on the table while varying the posture of their hand relative to the device. All participants wore eye masks during testing to eliminate visual input.

### Stimuli and stimulus framing

Tactile stimuli were produced with a custom-built device composed of two adjacent refreshable Braille cells forming a 4 × 4 pin array. Pins were activated sequentially to generate apparent rotational motion that traversed a roughly circular/tangential path across the stimulated skin area. The motion across the device was presented at 20 mm s^−1^ and lasted ≈ 1 s per trial.

We define device motion *d* ∈ {+1, −1 as +1} = clockwise (CW) and −1 = counterclockwise (CCW) when viewed from the device-facing (dorsal) side of the hand (device coordinates). Participants were instructed to judge motion direction relative to the local skin-tangent frame: report CW/CCW as if looking along the skin from the stimulated side. Responses were made with two keys (CW = +1, CCW = −1). On a subset of trials (see Supplementary Methods), participants also made a forced-choice parity judgment *p* ∈ {+1, −1} with *p* = +1 indicating the stimulus was perceived as originating from the skin-side, aka inward (skin vantage) and *p* = −1 indicating the stimulus was perceived as originating from the external or world-side/outward perspective (world vantage).

### Mappings and functional equations

Let *d* denote device motion in device coordinates. Let *R* denote the participant’s reporting-frame transform that maps device motion into the participant’s spatial reporting-frame motion sign; represent *R* as a function *R* : *D* → *S* where *D* = {+1, −1} and *S* = {+1, −1} . We model *R* with the function

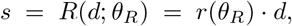

where *r*(*θ*_*R*_) ∈ {+1, −1} is a posture- and participant-dependent sign parameter determined by transform parameters *θ*_*R*_ (*r* = +1 preserves device sign; *r* = −1 inverts device sign).

Let *P* denote the participant’s parity-assignment function that maps the spatial/reporting-frame motion sign into a perceived vantage (skin vs. world). Define *P* : *S* → {+1, −1} with

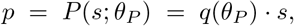

where *q*(*θ*_*P*_) ∈ {+1, −1} is a posture- and participant-dependent parity sign parameter determined by parameters *θ*_*P*_ (*q* = +1 indicates the reporting-frame orientation corresponds to a skin-side vantage; *q* = −1 indicates it corresponds to a world-side vantage).

Participants’ explicit reported motion sign *y* and parity *p* are therefore given by the composition of these mappings:

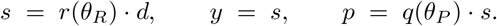

Equivalently,

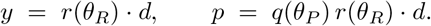

### Interpretation of parameter changes across postures

- If the reporting-frame sign *r*(*θ*_*R*_) changes sign across postures while *q*(*θ*_*P*_) remains constant, reported motion *y* will flip with posture but explicit parity *p* will remain stable. This pattern is consistent with a change in reporting frame (mapping from device to reporting coordinates) without a parity/vantage reassignment.
- If the parity sign *q*(*θ*_*P*_) changes sign across postures, then explicit parity *p* will flip; because *p* depends on *s*, flips in *p* that co-occur with flips in *y* indicate a true parity/vantage reassignment (skin ↔ world) rather than solely a reporting-frame sign change.
- Both *r*(*θ*_*R*_) and *q*(*θ*_*P*_) may vary across participants and skin sites (e.g., fingertip vs. palm). Visual experience may influence which parameter (or combination) is posture-sensitive.

### Procedure

In all three experiments, we used the method of single stimuli (26). Participants judged the direction of rotation (clockwise or counter-clockwise) relative to an internal standard, without receiving feedback. Prior to testing, participants were familiarized with clockwise and counter-clockwise motion using example stimuli. Each block comprised 16 clockwise and 16 counter-clockwise trials, presented in pseudorandom order. Randomised sequences were generated using seeded random initial values.

Analyses were designed to dissociate these alternatives by recording *y* and *p* on a subset of trials and testing whether posture-dependent flips in *y* co-occurred with flips in *p* (see Analysis). Our prediction was that if posture alters only the reporting-frame transform, we expect changes in *r*(*θ*_*R*_) with stable *q*(*θ*_*P*_); if posture induces parity reassignment, we expect joint changes in *r*(*θ*_*R*_) and *q*(*θ*_*P*_) such that *p* flips with *y*. Sighted and congenitally blind groups were hypothesized to differ in which parameter ( *r* or *q*) is posture-sensitive (15).

### Analysis

For Experiment 3, we assessed the orientation invariance of hand posture changes in blind participants’ surface assignment. The primary measure was the probability of surface interpretation, computed from 64 trials per condition: participant, posture, and motion direction. To evaluate whether participants exhibited greater consistency across orientation conditions than expected by chance, we conducted a nonparametric permutation test. For each participant, we computed the standard deviation of surface probability across four hand orientations. We then compared the observed group mean standard deviation to a null distribution generated by randomly shuffling orientation labels within each participant 10,000 times. The p-value was defined as the proportion of permuted samples with a mean standard deviation less than or equal to the observed value. This approach allowed us to determine whether the consistency in surface interpretation across orientations was statistically significant, independent of distributional assumptions and robust to individual idiosyncrasies.

## Supporting information

upplementary file 1. To assess whether tactile acuity across hand regions could be a factor in the results, we employed a kinetic version of the tacti

Supplementary file 2. To examine whether the stability of perceived motion direction across hand pos- ture and surface region (finger vs. palm) depen

## Acknowledgments

This work was supported by a Fyssen Foundation fellowship to J.H.O., a grant ANR-16-CE28-0015 “Developmental Tool Mastery” to V.H. and M.A., a Leverhulme Trust Visiting Professorship Grant to V.H.

## Additional information

## Author contributions

JHO, MA and VH conceived this work. JHO design and ran the experiments and analysed the data. JHO, MA and VH wrote the manuscript.

## Author ORCIDs

This research was supported by several organizations:

- The **Fyssen Foundation** provided a fellowship to J. Hartcher-O’Brien.
- The **Agence nationale de la recherche (ANR)** awarded grant number ANR-16-CE28-0015 to A. Farne.
- The **Leverhulme Trust** funded a visiting professorship (VP1-2016-060) for V. Hayward.

The funders had no role in study design, data collection, and interpretation, or the decision to submit the work for publication.

## Additional files

## Data availability

All data generated and analysed during this study will be made available at https://osf.io/

<u>Author(s) Year Dataset title Dataset URL Database and Identifier</u>

## Supplementary files

- Supplementary file 1. To assess whether tactile acuity across hand regions could be a factor in the results, we employed a kinetic version of the tactile “Landolt C-test” described in (27). This stimulus was like that shown in main text *Figure 1B* but with a gap that could be oriented in four possible directions. The hand posture was like in Experiment 1. The group of participants was the same as in Experiment 2. Half of the participants experienced clockwise stimuli while the other half experienced counterclockwise stimuli where a gap could appear at one of four possible locations in 80% of trials. There were 20% of the catch trials where the stimuli had no gap. The results, expressed in terms of sensitivity to stimulus orientation (*d*^*′*^ values), are shown in *Figure S1A*. Using affine regressions, tactile sensitivity failed to predict the participants’ responses, neither on the fingertip (*R*^2^ = 0.1, *F* (1, 15) = 0.47, *p >* 0.05), nor on the palm, (*R*^2^ = 0.3, *F* (1, 15) = 0.5, *p >* 0.05), see *Figure S1B*.
- Supplementary file 2. To examine whether the stability of perceived motion direction across hand posture and surface region (finger vs. palm) depended on stimulus direction, we decomposed responses by motion direction. Specifically, we calculated the proportion of trials in which participants reported feeling motion consistent with the actual stimulus direction‚Äîclockwise (CW) or counter-clockwise (CCW). That is, we assessed the proportion of CW-consistent responses for CW stimuli, and CCW-consistent responses for CCW stimuli, across all conditions. A repeated measures ANOVA revealed no significant effect of stimulus direction on the proportion of direction-consistent responses.

**Figure S1:**
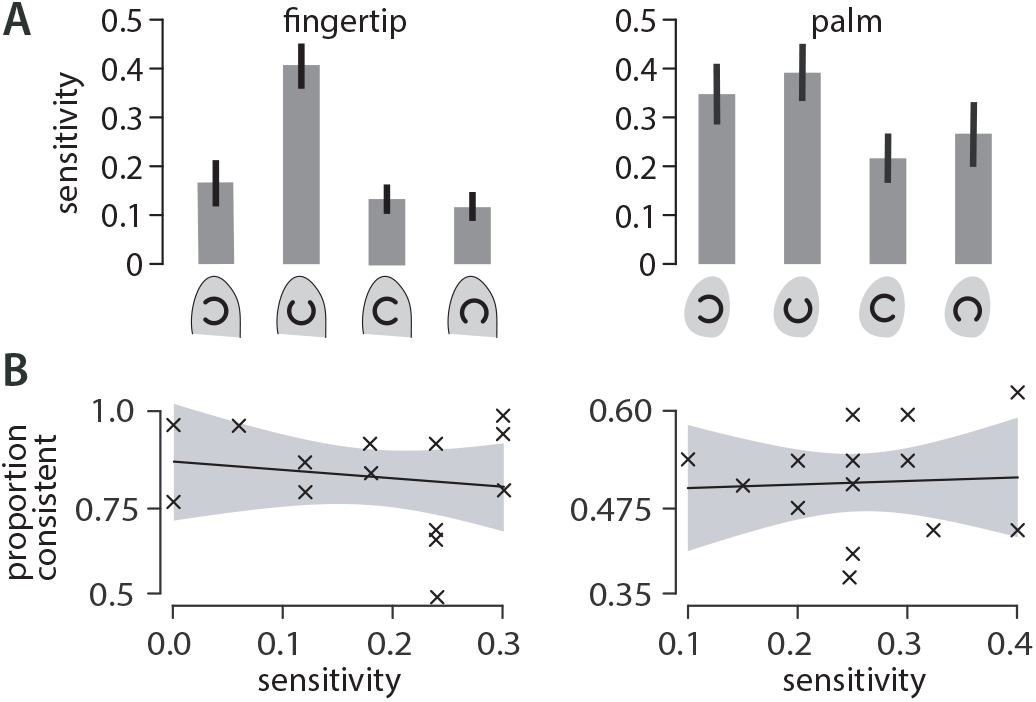
Sensitivity Experiment. (**A**) Detection sensitivity as a function of pattern orientation. Error bars show one standard deviation. **(B)** Affine regression of detection sensitivity against response type showed almost no correlation; 95% confidence regions shown in grey.

*Figure S2* shows the results from Experiment 2, separated by motion direction. The top panel shows responses to CW stimuli, and the bottom panel shows responses to CCW stimuli. Error bars represent one standard deviation.

**Figure S2:**
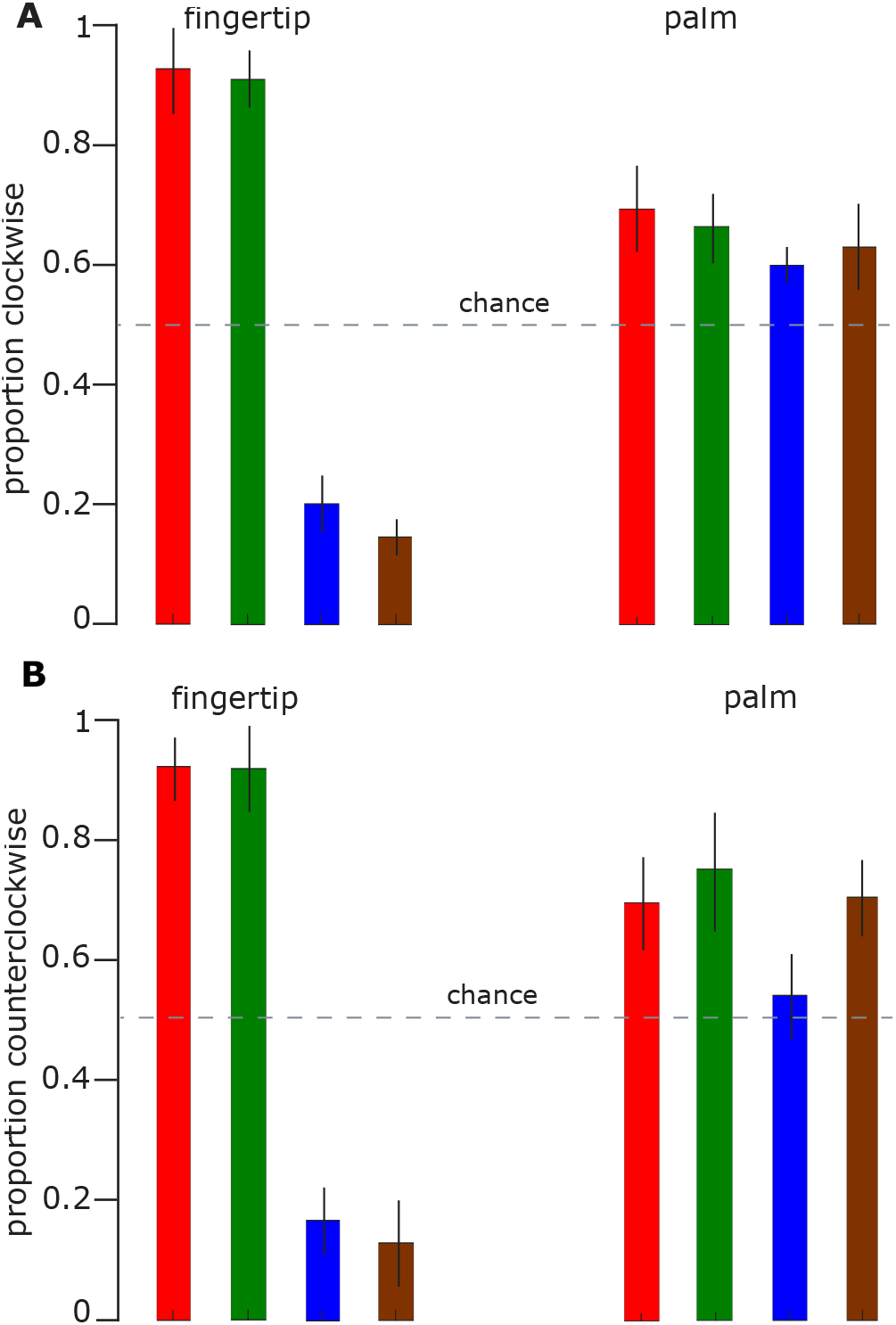
Experiment-2 non-collapsed motion direction responses for **(A)** Clockwise stimuli across the four postures, and **(B)** Counter-clockwise stimuli across the four postures tested. Error bars show one standard deviation.

## References

[1] Encyclopedia of Mathematics. https://encyclopediaofmath.org/wiki/Orientation, 2010.

[2] F. N. Newell, M. O. Ernst, B. S. Tjan, and H. H. Bülthoff. Viewpoint dependence in visual and haptic object recognition. Psychological Science, 12(1): 37–42, 2001.

[3] Tamar R Makin, Alona O Cramer, Jan Scholz, Avital Hahamy, David Henderson Slater, Irene Tracey, and Heidi Johansen-Berg. Deprivation-related and use-dependent plasticity go hand in hand. Elife, 2:e01273, 2013.

[4] Luigi Tamè, Andrew Moles, and Nicholas P Holmes. Within, but not between hands interactions in vibrotactile detection thresholds reflect somatosensory receptive field organization. Frontiers in psychology, 5:174, 2014.

[5] Nicholas P Holmes. Hand-centered space, hand-centered attention, and the control of movement. 2013.

[6] G. Arnold and M. Auvray. Perceptual learning: Tactile letter recognition transfers across body surfaces. Multisensory Research, 46(29–38), 2014.

[7] G. Arnold, C. Spence, and M. Auvray. Taking someone else’s spatial perspective: Natural stance or effortful decentring? Cognition, 148: 27–33, 2016.

[8] E. Azañón, M. R. Longo, S. Soto-Faraco, and P. Haggard. The posterior parietal cortex remaps touch into external space. Current Biology, 20: 1304–1309, 2010.

[9] E. Azañón and S. Soto-Faraco. Changing reference frames during the encoding of tactile events. Current Biology, 18: 1044–1049, 2008.

[10] Geza Révész. Experimental study in abstraction in monkeys. Journal of Comparative Psychology, 5(4): 293, 1925.

[11] Lynette A Jones and Susan J Lederman. Human hand function. Oxford university press, 2006.

[12] Jennifer Marie Groh. Coordinate transformations, sensorimotor integration and the neural basis of saccades to somatosensory targets. University of Pennsylvania, 1993.

[13] Anton R Sobinov and Sliman J Bensmaia. The neural mechanisms of manual dexterity. Nature Reviews Neuroscience, 22(12): 741–757, 2021.

[14] Elena Azañón, Kim Mihaljevic, and Matthew R. Longo. A three-dimensional spatial characterization of the crossed-hands deficit. Cognition, 157: 289–295, 2016.

[15] Xavier Job, Gabriel Arnold, Louise P Kirsch, and Malika Auvray. Vision shapes tactile spatial perspective taking. Journal of Experimental Psychology: General, 150(9): 1918, 2021.

[16] Xavier E Job, Louise P Kirsch, and Malika Auvray. Spatial perspective-taking: insights from sensory impairments. Experimental brain research, 240(1): 27–37, 2022.

[17] T. Natsoulas and R. A. Dubanoski. Inferring the locus and orientation of the perceiver from responses to stimulation of the skin. The American journal of psychology, 77(2): 281–285, 1964.

[18] Jerome E. Podell. Ontogeny of the locus and orientation of the perceiver. Child Development, pages 993–997, 1966.

[19] D. W. Corcoran. The phenomena of the disembodied eye or is it a matter of personal geography? Perception, 1977.

[20] L. M. Parsons and S. Shimojo. Perceived spatial organization of cutaneous patterns on surfaces of the human body in various positions. Journal of Experimental Psychology: Human Perception and Performance, 13(3): 488, 1987.

[21] S. Shimojo, M. Sasaki, L. M. Parsons, and S. Torii. Mirror reversal by blind subjects in cutaneous perception and motor production of letters and numbers. Perception & Psychophysics, 45: 145–152, 1989.

[22] J. Hartcher-O’Brien and M. Auvray. Cognition overrides orientation dependence in tactile viewpoint selection. Experimental Brain Research, 234(7): 1885–1892, 2016.

[23] E. R. Ferrè, C. Lopez, and P. Haggard. Anchoring the self to the body: vestibular contribution to the sense of self. Psychological Science, 25(11): 2106–2108, 2014.

[24] L. Dupin, V. Hayward, and M. Wexler. Radial trunk-centred reference frame in haptic perception. Scientific Reports, 8:13550, 2018.

[25] O. Gapenne, K. Rovira, A. Ali Ammar, and C. Lenay. Tactos: Special computer interface for the reading and writing of 2d forms in blind people. In Universal Access in HCI, Inclusive Design in the Information Society, volume 10, pages 1270–1274. CRC Press, 2003.

[26] S. P. McKee, G. H. Silverman, and K. Nakayama. Precise velocity discrimination despite random variations in temporal frequency and contrast. Vision Research, 26(4): 609–619, 1986.

[27] D. Chan. An apparatus for the measurement of tactile acuity. The American Journal of Psychology, 77(3): 489–491, 1964.

