## Supplementary figures and images for "Inside Out: Functional Flexibility in Assigning the Body’s inside-outside"

### Supplementary file 2. To examine whether the stability of perceived motion direction across hand pos- ture and surface region (finger vs. palm) depen

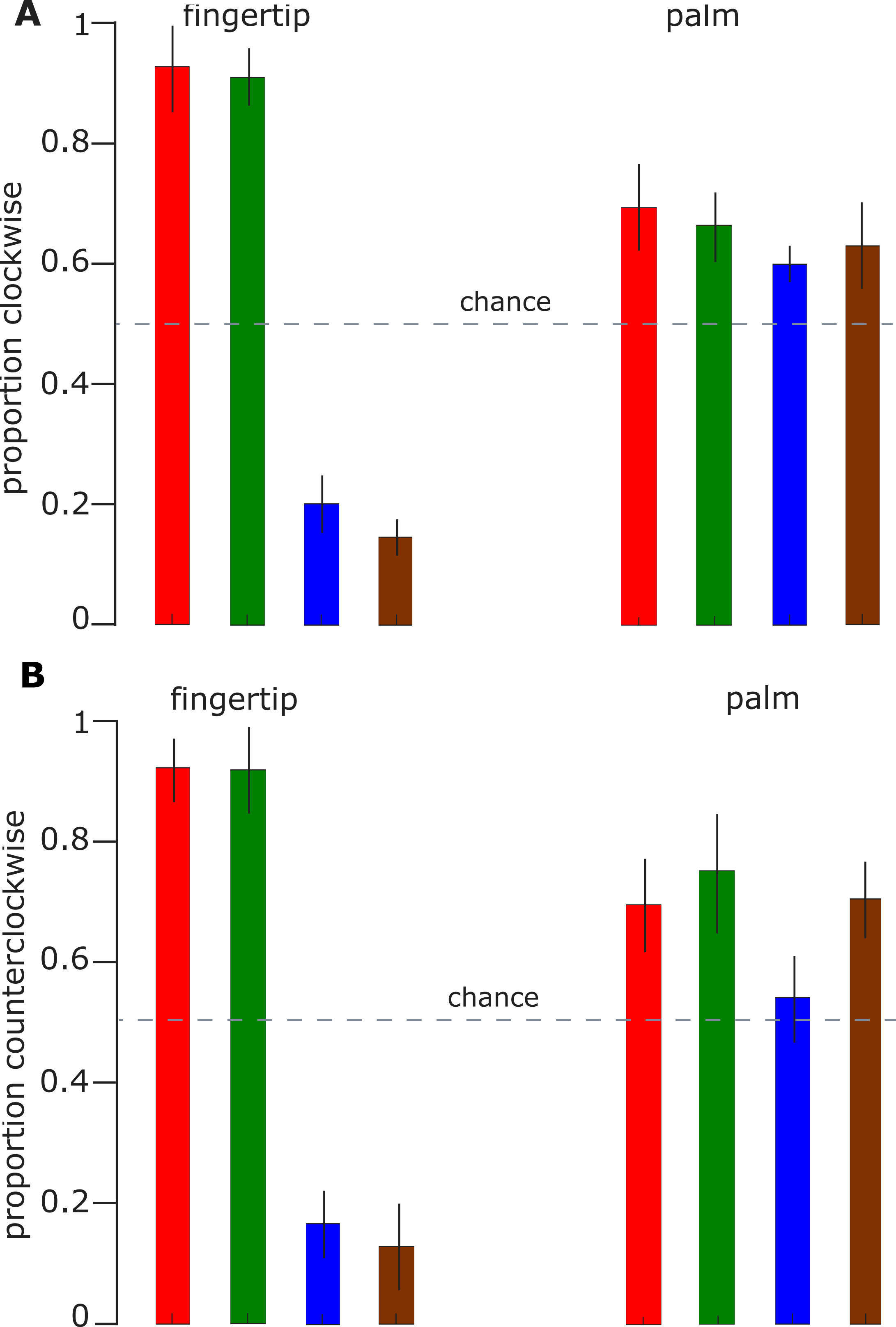

### upplementary file 1. To assess whether tactile acuity across hand regions could be a factor in the results, we employed a kinetic version of the tacti

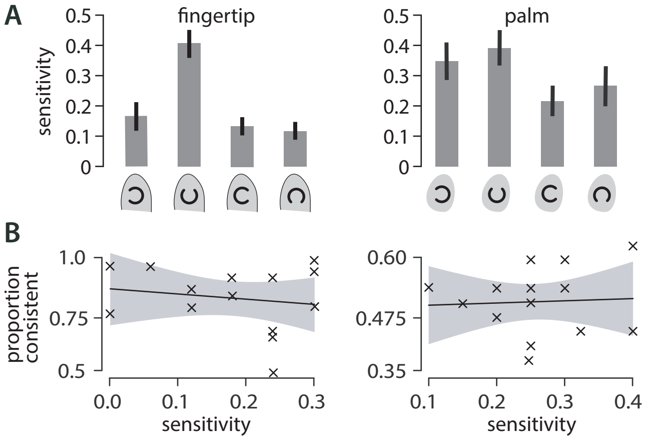
